# The co-stimulatory activity of Tim-3 requires Akt and MAPK signaling and immune synapse recruitment

**DOI:** 10.1101/2019.12.30.878520

**Authors:** Shunsuke Kataoka, Priyanka Manandhar, Judong Lee, Creg J. Workman, Hridesh Banerjee, Andrea L. Szymczak-Workman, Michael Kvorjak, Jason Lohmueller, Lawrence P. Kane

**Affiliations:** Department of Immunology University of Pittsburgh Pittsburgh, PA 15261; Asahi-Kasei Pharma Corporation Shizuoka, Japan; Graduate Program in Microbiology and Immunology University of Pittsburgh Pittsburgh, PA 15261; Department of Surgery, Hillman Cancer Center, University of Pittsburgh, Pittsburgh, PA 15261

## Abstract

Expression of the transmembrane protein Tim-3 is increased on dysregulated T cells undergoing chronic T cell activation, including in chronic infection and solid tumors. We and others previously reported that Tim-3 exerts apparently paradoxical co-stimulatory activity in T cells (and other cells), including enhancement of ribosomal S6 protein phosphorylation (pS6). Here we examined the upstream signaling pathways that control Tim3-mediated increases in pS6 in T cells. We have also defined the localization of Tim-3 relative to the T cell immune synapse and impacts on downstream signaling. Recruitment of Tim-3 to the immune synapse was mediated exclusively by the transmembrane domain, replacement of which impaired Tim-3 co-stimulation of pS6. Strikingly, enforced localization of the Tim-3 cytoplasmic domain to the immune synapse in the context of a chimeric antigen receptor still allowed for robust T cell activation. Our findings are consistent with a model whereby Tim-3 enhances TCR-proximal signaling under acute conditions.

**One Sentence Summary:** Here we define elements of signaling and localization associated with Tim-3 co-stimulatory function in T cells.

## Introduction

T cell activation and effector function are orchestrated by a complex series of cell-cell interactions between T cells and antigen presenting cells (APCs). The most important interaction is between T cell receptor for antigen (TCR) and major histocompatibility complex (MHC) proteins, which present peptide antigens to the T cell. Recognition of peptide/MHC by a TCR is translated into biochemical signaling events by the TCR-associated CD3 γ, δ and ε and ζ signaling chains (*1, 2*). While these events are critical for T cell activation, there are numerous other receptor-ligand interactions that also regulate – either positively or negatively – the ultimate effects on T cell activation and function. The negative regulators in particular have garnered much attention in recent years, and blocking antibodies to some of these (CTLA-4 and PD-1) are now clinically approved for use in selected solid tumors (*3, 4*). In a parallel series of studies, the use of engineered T cells expressing chimeric antigen receptors (CARs) directed at specific tumor antigens has gained substantial traction in the treatment of specific hematological malignancies (*5*). The efficacy of the latter approach is also affected by both the expression of endogenous positive and negative regulators in the CAR T cells, as well as the inclusion of different co-stimulatory signaling domains in the CAR itself.

The protein T cell (or transmembrane) immunoglobulin and mucin domain 3 (Tim-3) was originally described as a marker for Th1 helper T cells (*6*). Subsequent studies revealed that Tim-3 is also expressed on acutely activated CD8^+^ T cells and a subset of regulatory T cells, in addition to various non-T cells, including some macrophages, dendritic cells, NK cells and mast cells (*7–9*). However, most of the attention paid to Tim-3 in recent years has focused on the high expression of Tim-3 on “exhausted” CD8^+^ T cells observed under conditions of chronic infection or within the tumor microenvironment. These cells comprise a subset of the cells that express the immune checkpoint molecule PD-1, which was previously described as the most robust marker for exhausted T cells; indeed, monoclonal antibodies (mAb’s) that interfere with PD-1 interactions with its ligands (PD-L1, PD-L2) have demonstrated efficacy in a subset of cancer patients. Furthermore, T cells expressing high levels of both PD-1 and Tim-3 appear to be even more dysfunctional than those expressing PD-1 alone, as originally shown in chronic viral infection (*10–13*). Consistent with this notion, dual blockade of PD-1 and Tim-3 has a modestly enhanced ability to “rescue” the function of a population of exhausted T cells (*10, 11, 14*). Several antibodies directed against Tim-3 are now in clinical trials and combination therapies are being actively explored (*15*). Thus, a prominent model that has emerged is that Tim-3 primarily functions as a negative regulator of T cell activation, in a manner similar to PD-1. Recent correlative evidence for this model was provided by the discovery of patients bearing germline loss-of-function mutations in Tim-3 (*16*).

Despite these data supporting the notion that Tim-3 is a dominant negative regulator, a number of observations complicate such a straightforward model. First, there has yet to emerge a clear-cut mechanism by which Tim-3 might transmit an inhibitory signal, for example through recruitment of phosphatases like PD-1, or by competition with co-stimulatory receptors, like CTLA-4. Second, at least four ligands have been described to interact with Tim-3, including galectin-9, phosphatidylserine (PS), HMGB1 and CEACAM1 (*7–9*), which are known to also interact with other and distinct receptors (*15, 17–20*), making the definition of important receptor-ligand interactions difficult at best. Third, studies from multiple labs on both T cells and non-T cells have directly or indirectly demonstrated an ability of Tim-3 to transmit positive, activation-promoting, signals (*21–26*). Our own work has suggested a model by which expression of Tim-3 might either promote the development of T cell exhaustion by enhancing or sustaining TCR signaling or actually represent a “last-ditch” effort by exhausted T cells to maintain some function (*22, 27*). Indeed, recent mechanistic studies on PD-1 blockade have demonstrated that the main target of this therapy is a pool of Tcf1-expressing “stem-like” CD8^+^ T cells that express intermediate levels of PD-1 but no Tim-3 (*28–31*), while the few cells that express Tim-3 alone are actually capable of robust proliferation (*32*). In addition, tumor-infiltrating CD8^+^ T cells expressing Tim-3 and high levels of PD-1 have been demonstrated to adopt a distinct epigenetic program that serves as a barrier to rejuvenation by checkpoint blockade (*33*).

A previous study provided evidence that Tim-3 is recruited to the immune synapse (IS)(*34*), a structure that forms between T cells and antigen presenting cells during productive antigen recognition, using standard confocal microscopy. In addition, this study suggested that IS recruitment of Tim-3 contributed to its negative regulatory function, although the precise mechanism was not defined. Here, we have examined the relationship between Tim-3 and the IS using both imaging cytometry with T cell-APC conjugates and total interference reflection fluorescence (TIRF) microscopy with lipid bilayers. We report that there is robust recruitment of Tim-3 to the IS, which lags recruitment of the TCR/CD3 complex to the IS. Furthermore, we demonstrate that IS recruitment of Tim-3 is mainly mediated by its transmembrane (TM) domain. By substituting the TM, we show that IS recruitment of Tim-3 correlates with the previously reported co-stimulatory activity of the protein. Finally, we show that even under conditions of enforced IS recruitment with a chimeric antigen receptor (CAR), the cytoplasmic domain of Tim-3 provides a co-stimulatory, but not inhibitory, signal for T cell activation. Thus, our data are consistent with a model whereby the primary proximal effect of Tim-3 signal is to support T cell activation.

## Results

### Enhancement of S6 phosphorylation by Tim-3 is mediated by both MEK/ERK and PI3K-Akt

We previously showed that ectopic expression of Tim-3 in T cells can enhance phosphorylation of ribosomal protein S6 (pS6), in conjunction with TCR signaling (*22, 24*). S6 phosphorylation can be promoted by activation of MEK/ERK and/or PI3K/Akt signaling (*35*). To assess whether Tim-3-mediated ERK and/or Akt activation are associated with the enhancement of pS6, we used a highly selective inhibitor of MEK1 and MEK2 (U0126) or an allosteric inhibitor of Akt (Akt*i*1/2). We confirmed that U0126 and Akt*i*1/2 significantly blocked pERK and pAkt, respectively, in a dose-dependent manner (**fig. S1A**). We next co-transfected Jurkat T cells with a GFP-encoding plasmid (as a control for transfection) and either empty vector or Flag-tagged Tim-3. We then stimulated these cells with αTCR mAb, with or without inhibitors, and detected Tim-3 expression with α-Flag antibody (**Fig. 1A**). As we reported previously, Tim-3 expression enhanced TCR-induced pS6 in the transfected (GFP+) population. We also observed that either U0126 or Akt*i*1/2 treatment inhibited pS6 induction, although neither was sufficient to completely block the enhancement of pS6 by Tim-3 expression. However, combined treatment with both inhibitors reduced pS6 more than either inhibitor alone and repressed the co-stimulatory activity of Tim-3, indicating that both MEK/ERK and PI3K-Akt pathways play a role in the regulation of pS6 by Tim-3 expression (**Fig. 1B-C**).

**Figure 1.**
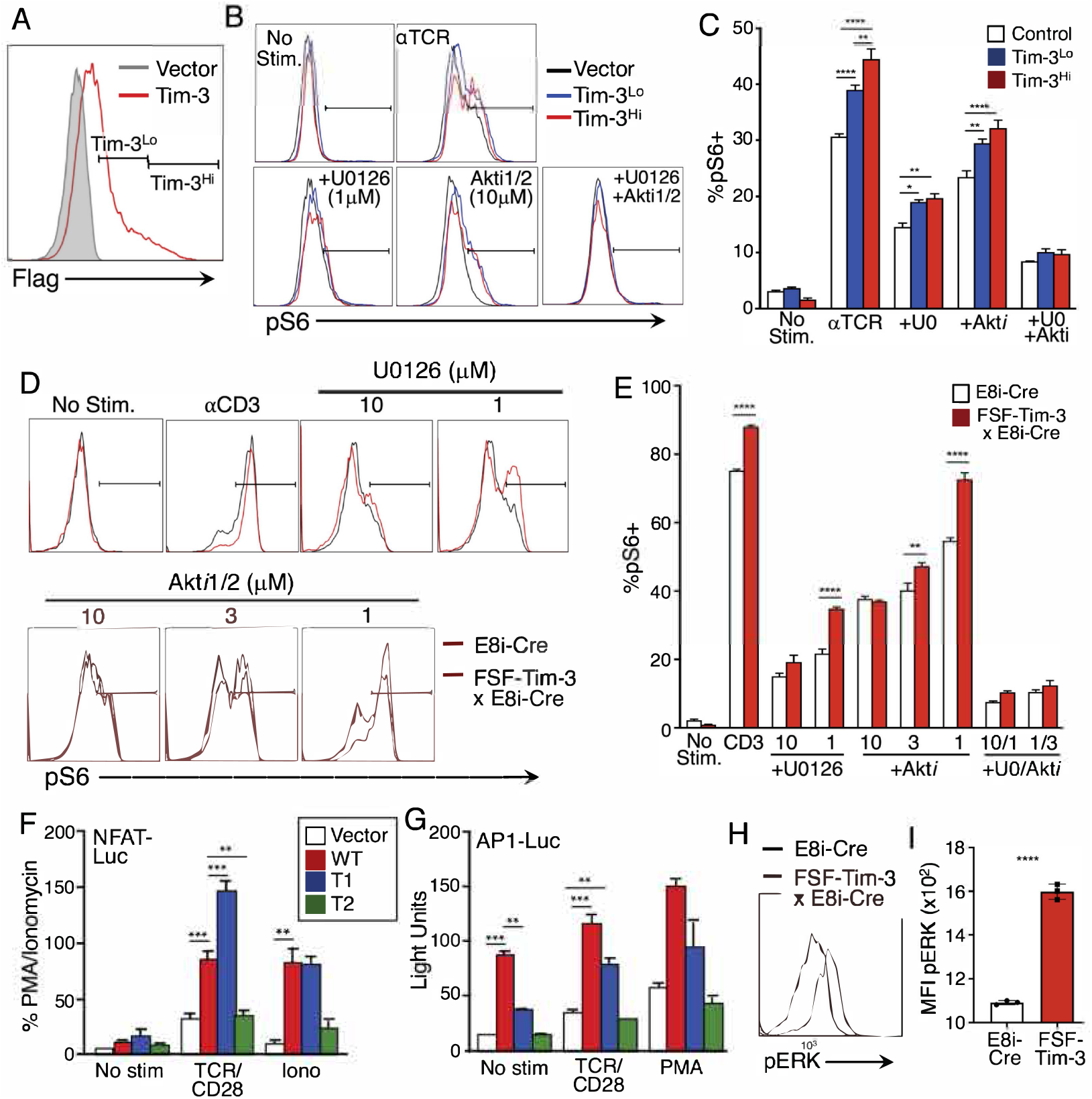
Tim-3 enhancement of pS6 requires both MEK-ERK and PI3K-Akt signaling. (**A**) Expression of Flag-tagged Tim-3 on transfected Jurkat T cells shown by flow cytometry. (**B**) Jurkat T cells were pretreated with vehicle, U0126, or Akt*i*1/2 before being stimulated with αTCR for 30 mins. pS6 (Ser^236/236^) was analyzed by flow cytometry, within control, Tim-3^Lo^ or Tim-3^Hi^ cells. (**C**) Quantification of %pS6^+^ cells based on flow cytometry shown in B. (**D**) Primary T cells were pretreated with vehicle, U0126, or Akt*i*1/2 before being stimulated with αCD3 for 4 hrs. pS6 was analyzed within CD8^+^ T cells from E8i-Cre or E8i-Cre/FSF-Tim-3 mice. (**E**) Quantification of %pS6^+^ cells as shown in D. (**F-G**) Luciferase reporter assays on Jurkat T cells transfected with vector control or the indicated Tim-3 constructs plus either NFAT-luciferase (F) or AP-1-luciferase (G) reporter construct. Cells were stimulated as indicated and luciferase activity was quantified in a luminometer. (**H-I**) Induction of pERK in WT (E8i-Cre only) or Tim-3 Tg T cells. Representative flow cytometry histograms are shown in (H) and aggregated quantified data of three independent samples are shown in (I). Data in each panel are representative of at least three experiments. Data shown in the bar graphs represent the mean ± SEM. \**P* < 0.05, \*\**P* < 0.01, \*\*\*\**P* < 0.0001, Tukey’s multiple comparisons test of triplicate samples. Data in panels A-C are representative of seven experiments each; D-I are representative of three experiments.

Jurkat T cells lack expression of the PTEN phosphatase (*36*), rendering these cells hyper-responsive to TCR stimulation. To confirm the above findings in primary T cells, we used CD8-specific inducible Tim-3 mice (**fig. S1B**). These mice were obtained by crossing transgenic *Rosa26* knock-in “flox-stop-flox” Tim-3 (FSF-Tim-3) mice with CD8 T cell-specific E8i-Cre mice (*22*). Naïve T cells from E8i-Cre and E8i-Cre x FSF-Tim-3 mice were pretreated with the above-described inhibitors before being stimulated through the TCR. Consistent with our previous findings (*22*), pS6 was enhanced in CD8^+^ T cells from E8i-Cre x FSF-Tim-3 mice (**Fig. 1D-E**). Treatment with either inhibitor at high concentrations was sufficient to block the enhancement of pS6 by Tim-3, whereas combined treatment with moderate doses of both inhibitors resulted in a decrease of pS6 more than either inhibitor alone. We also confirmed the effects of these inhibitors on pS6 in the murine D10 T cell line (*37*). Consistent with the above results, combined treatment with both inhibitors potently blocked induction of pS6 (**fig. S1C-E**). These results suggest that co-stimulation of S6 protein phosphorylation by Tim-3 requires both the MEK-ERK and PI3K-Akt pathways to enhance TCR signaling.

To further analyze the contribution of Tim-3 to activation of MEK-ERK, we transfected Jurkat T cells with luciferase reporter constructs driven by NFAT or AP-1 elements, which are activated by calcium or Ras/MAPK signaling, respectively (*38*). We also transfected these cells with WT or truncated forms of Tim-3 (**Fig. S2A**). Thus, as we previously showed with a composite NFAT/AP-1 reporter (*24*), WT Tim-3 and a construct with a partially truncated cytoplasmic tail (T1) efficiently co-stimulated the activation of an NFAT-driven reporter, in the presence of TCR/CD28 or ionomycin stimulation (**Fig. 1F**). By contrast a Tim-3 construct with a more severe cytoplasmic tail turncation did not co-stimulate NFAT-dependent transcription. We also assessed the effects of the same Tim-3 constructs on an AP-1 reporter. Unlike the results obtained with NFAT/AP-1 or NFAT reporters, a pure AP-1 reporter was activated by co-transfection with WT Tim-3 alone, without a need for additional stimulation (**Fig. 1G**). This suggested an ability of Tim-3 to activate upstream MEK-ERK signaling in the basal state. We addressed this further by examining ERK phosphorylation directly in the FSF-Tim-3 mice described above. Thus, CD8+ T cells expressing transgenic Tim-3 displayed higher basal phosphorylation of ERK than control E8i-Cre T cells (**Fig. 1H-I**). These results demonstrate that Tim-3 has an intrinsic ability to activate MEK-ERK signaling, while its co-stimulatory activity may function more through the calcium-NFAT pathway.

### Amnis ImageStream analysis reveals Tim-3 recruitment to the IS

Co-stimulatory and co-inhibitory receptors often colocalize with TCR at the immunological synapse (IS) (*39*). A previous study suggested that Tim-3 is recruited to the IS in activated T cells (*34*). To better understand the mechanisms underlying co-stimulation by Tim-3 in T cells, we examined Tim-3 localization during IS formation on Jurkat T cells transfected with Flag-tagged Tim-3, using the Amnis ImageStream flow cytometry system. Raji cells (a human B cell lymphoma line) pulsed with SEE were used as APCs, and IS formation was confirmed on the basis of CD3 recruitment to the Jurkat T cell-Raji cell interface (**Fig. 2A-B**). We did observe an increase in the proportion of Tim-3 recruitment to the IS over time, although Tim-3 displayed a lower overall degree of recruitment, with somewhat slower kinetics, compared with the behavior of CD3. To determine whether the same pattern of recruitment of Tim-3 is also seen in primary murine T cells, we employed P14 TCR transgenic mice, with or without expression of Tim-3 from a *Rosa26* knock-in cassette (*22*). CD8^+^ T cells from these mice were stimulated with autologous T cell-depleted splenocytes, with or without the cognate peptide antigen, the gp33 peptide from LCMV. Thus, consistent with the results obtained with Jurkat T cells, we observed enhanced recruitment of Tim-3 to the IS of CD8^+^ P14 TCR Tg x Tim-3 Tg T cells in the presence of cognate peptide (**Fig. 2C-D**).

**Figure 2.**
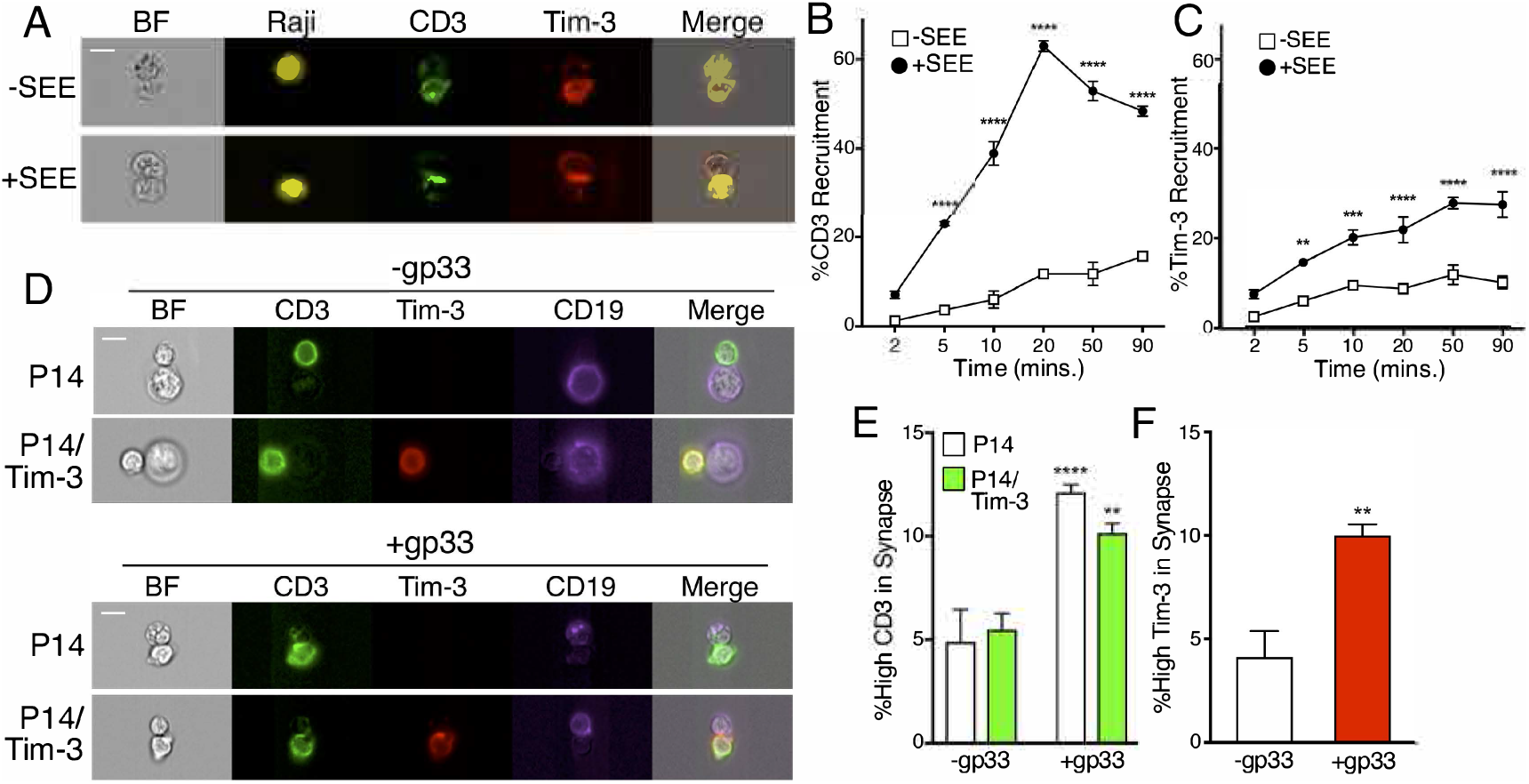
Recruitment of Tim-3 to the immune synapse of Jurkat and P14 Tg T cells. (**A**) Jurkat T cells transfected with Flag-tagged Tim-3 (red) were mixed with Raji cells (yellow), either pulsed with SEE or unpulsed, at a ratio of 1:1 for 20 mins. and then were analyzed by ImageStream. (**B-C**) Conjugates were incubated for the indicated times. The percentage of either CD3 or Tim-3 recruitment was quantified from the ratio of the intensity of either CD3 (green) or Tim-3 (red) within the interface mask of conjugates containing one Jurkat T cell and Raji cell to that of the whole cell mask, using IDEAS software. Each point represents an average of 291 conjugates. (**D-F**) CD8^+^ T cells from control P14 TCR Tg mice or those expressing Cre-inducible Tim-3 were mixed with purified B cells from the same mice, with or without gp33 peptide, and conjugates were analyzed as in panels B-C. Each point represents an average of 418 conjugates in the absence of antigen and 5500 cells in the presence of antigen. The scale bars in all panels indicate 10μM. Data in panels A-C are representative of two experiments; D-F are representative of four experiments.

### The extracellular and intracellular domains of Tim-3 are dispensable for its recruitment to the immune synapse

We next sought to determine the roles of specific Tim-3 domains in its recruitment to the IS. Using various mutants of Tim-3, we previously showed that the cytoplasmic tail of Tim-3 is required for its co-stimulatory activity, at least at the level of anti-TCR-induced activation of NFAT/AP-1 and NF-κB transcriptional reporters (*24*). To address the relationship between Tim-3 signaling and its localization, we used Flag-tagged Tim-3 variants lacking either the sequence encompassing three tyrosines at positions 271, 272 and 274 (truncation 1; T1) or a more severely truncated variant also lacking tyrosines 256 and 263 (truncation 2; T2) (**fig. S2A**). We previously showed that the T1 construct retains Tim-3 co-stimulation of NFAT/AP-1 activity, while the T2 construct abolishes this activity (*24*). Consistent with those data, we found that the T1 mutant still retained the ability to enhance pS6 comparably to WT Tim-3, whereas the T2 truncation abolished the ability of Tim-3 to enhance pS6 (**fig. S2E-F**). Surprisingly, however, we noted no difference in the localization of these constructs to the IS upon stimulation with SEE-pulsed Raji cells, compared to full-length Tim-3 (**Fig. 3A-C**). The T2 construct still retains 38 amino acids downstream of the transmembrane domain. To more rigorously address the role of the intracellular domain in Tim-3 IS recruitment, we used a Flag-tagged Tim-3 variant lacking the entire cytoplasmic region (ΔCyto; **fig. S2A**). However, there was no significant difference in the localization of the ΔCyto construct, compared to full-length Tim-3 (**Fig. 3D-F**). Thus, IS recruitment of Tim-3 does not require its cytoplasmic domain.

**Figure 3.**
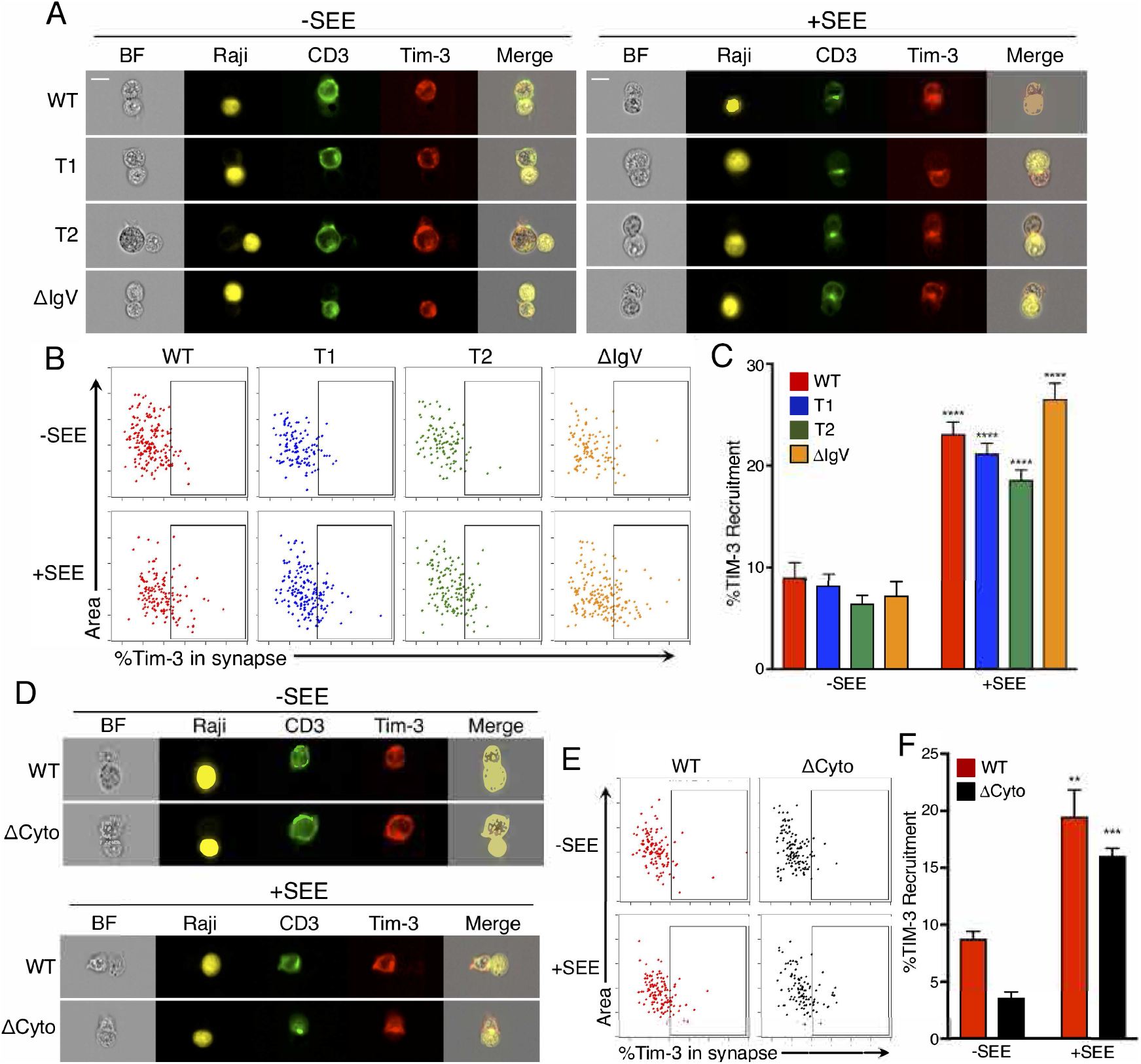
The IgV domain and cytoplasmic tail are dispensable for Tim-3 recruitment to the IS. **(A)**Representative images of Jurkat T cells transfected with either Flag-tagged WT Tim-3 or partial cytoplasmic truncations (T1, T2) or ΔIgV constructs (red) and mixed with Raji cells (yellow), either pulsed with SEE or unpulsed, at a ratio of 1:1 for 20 mins. (**B**) Representative dot plots analyzing the ratio of the intensity of WT or mutant Tim-3 constructs in the interface mask between a transfected Jurkat T cell and Raji cell to that of the whole cell mask. (**C**) Quantification of %Tim-3 recruitment. Each bar represents an average of 147 conjugates. (**D**) Representative images of Jurkat T cells transfected with either Flag-tagged WT or ΔCyto Tim-3 (red) and mixed with Raji cells (yellow), either pulsed with SEE or unpulsed, at a ratio of 1:1 for 20 mins. (**E**) Representative dot plots analyzing the ratio of the intensity of WT or ΔCyto Tim-3 in the interface mask between a transfected Jurkat T cell and Raji cell to that of the whole cell mask. (**F**) Quantification of %Tim-3 recruitment. Each bar represents an average of 71 cells. Data represent the mean ± SEM. \*\**P* < 0.01, \*\*\**P* < 0.001 vs. −SEE, Tukey’s multiple comparisons test. The scale bars in all panels indicate 10μM. Experiments comparing WT, T1 and T2 constructs were performed at least three times; those with ΔCyto and ΔIgV were performed twice.

Previous studies have shown that receptor-ligand binding, e.g. B7 to CD28, can regulate the recruitment of immune co-stimulatory and inhibitory molecules to the IS (*40, 41*). The Tim-3 IgV domain has been shown to bind to various and diverse ligands such as galectin-9, PS, HMGB1 and Ceacam-1 (*42*) (*43–45*), which constitute all known Tim-3 ligands. We therefore used a Flag-tagged Tim-3 variant lacking the IgV domain (ΔIgV) (**fig. S2A**), to address the role of ligand binding in Tim-3 IS recruitment. As with the cytoplasmic tail deletions, we observed that deletion of the IgV domain did not impair Tim-3 recruitment (**Fig. 3A-C**), indicating that Tim-3 recruitment to the IS may not require ligand binding, since deletion of the IgV domain eliminates the ability of Tim-3 to bind to all of its known ligands.

### The transmembrane domain of Tim-3 regulates its co-stimulatory activity and recruitment to the immune synapse

Previous studies suggested that the transmembrane domain of some cell-surface molecules can regulate recruitment to lipid rafts, cholesterol-rich membrane sub-regions that accumulate at the IS. We examined the role of the transmembrane domain of Tim-3 in IS recruitment, using a Tim-3-CD71 transmembrane chimeric protein (CD71tm), in which we replaced the transmembrane domain of Tim-3 with the corresponding domain of CD71. The CD71 TM domain has been shown to mediate exclusion from lipid rafts in the context of a chimeric receptor (*46*). Indeed, we observed a decrease in Tim-3 localization at the IS upon stimulation with SEE-pulsed Raji cells, in Jurkat T cells transfected with CD71tm Tim-3, compared with WT Tim-3 (**Fig. 4A-C**). Thus, the transmembrane domain of Tim-3 is critical for its localization to the IS.

**Figure 4.**
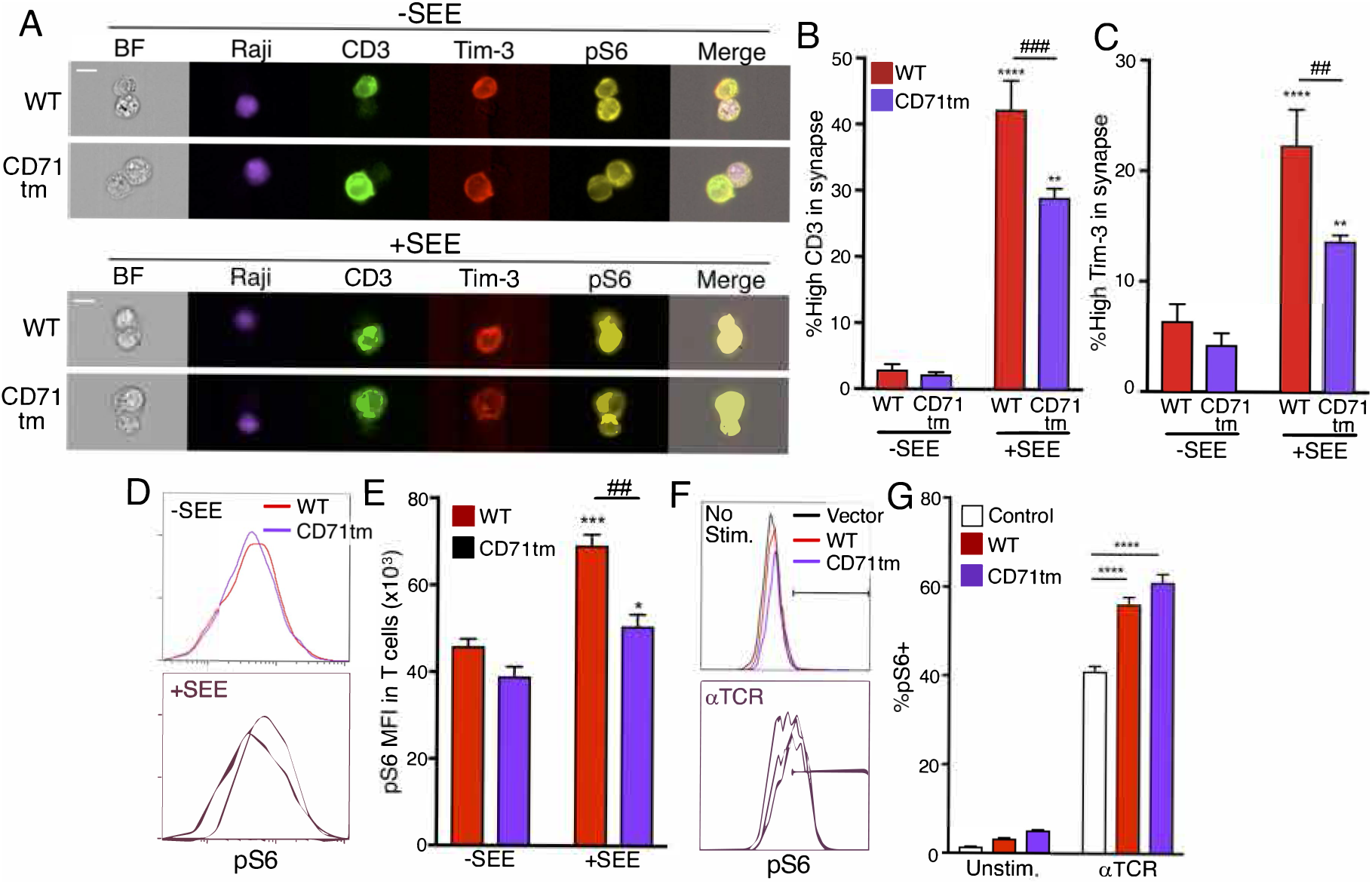
Role of the transmembrane domain in Tim-3 localization and function during IS formation. (**A**) Representative images of pS6 (yellow) in Jurkat T cells transfected with either Flag-tagged WT or CD71tm Tim-3 (red) and mixed with Raji cells (purple) either pulsed with SEE or unpulsed, at a ratio of 1:1, for 20 mins. (**B-C**) Quantification of CD3 (**B**) and Tim-3 (**C**) enrichment in the synapses of multiple Jurkat-Raji conjugates, with or without SEE. Each bar represents 85 conjugtaes without SEE and 203 conjugates with SEE. (**D**) Representative histogram analyzing pS6 intensity in transfected Jurkat T cells in conjugates, with or without SEE. (**E**) Quantification of pS6 MFI across multiple conjugates. (**F**) pS6 was analyzed by flow cytometry, within control, WT Tim-3 or Tim-3 CD71tm-transfected Jurkat T cells, stimulated with αTCR for 30 mins. (**G**) quantification of %pS6^+^ cells. Each panel is representative of at least two independent experiments. Data shown represent the mean ± SEM. \**P* < 0.05, \*\*\**P* < 0.001, \*\*\*\**P* < 0.0001 vs. −SEE, ^##^*P* < 0.01 Tukey’s multiple comparisons test. The scale bars in all panels indicate 10μM. Data in panesl A-C are representative of three experiments; D-G are represenative of two experiments.

To assess the role of the transmembrane domain of Tim-3 in T cell activation during IS formation, we stimulated transfected Jurkat T cells with Raji cells pulsed with SEE. Thus, we observed that Jurkat T cells transfected with WT Tim-3 showed higher pS6 levels than cells transfected with CD71tm Tim-3, after conjugation with SEE-pulsed Raji cells (**Fig. 4D-E**). To assess whether replacement of the transmembrane domain of Tim-3 affects T cell activation more generally, we stimulated Jurkat T cells expressing either Flag-tagged WT Tim-3 or CD71tm Tim-3 with α-TCR mAb. However, in this context, we found similar pS6 levels between cells expressing WT and CD71tm forms of Tim-3 (**Fig. 4F-G**), indicating that CD71tm Tim-3 retains the ability to enhance pS6 after antibody stimulation, which does not result in IS formation. These data suggest that IS localization of Tim-3 is required for the co-stimulatory effect of Tim-3 on T cell activation, specifically under conditions inducing IS formation.

In the above experiments, we assessed the localization of Tim-3 and formation of immune synapse between T cells and APCs using imaging cytometry. This approach has the advantages of visualizing IS formation in a native T cell-APC setting as well as being amenable to high-throughput data collection. Nonetheless, we wanted to also assess Tim-3 localization in a system that allows for imaging of these structures at higher resolution and in real-time. We therefore chose the lipid bilayer system described by Dustin and colleagues (*47*). We paired this with TIRF microscopy and a fluorescence activating protein (FAP) chimeric approach, which together allow for high signal:noise imaging of protein localization at the plasma membrane (*48*). Primary murine T cells were activated in vitro and transduced with lentivirus encoding WT or CD71tm Tim3-FAP fusion proteins (see schematic in **Fig. 5A**). Representative expression of the constructs in the packaging cells and transduced T cells are shown (**fig. S3B-C**). Transduced cells were sorted and imaged on lipid bilayers containing ICAM-1 and anti-TCR beta Fab fragment. Consistent with results shown above, we observed recruitment of WT Tim-3 to the IS, as defined by TCR/CD3 recruitment (**Fig. 5B**). We next imaged the CD71tm variant of Tim-3 described above, using the same system. Again, consistent with results obtained using APC conjugates, the CD71tm Tim-3 construct was recruited less efficiently to the IS formed on lipid bilayers (**Fig. 5C**). Most strikingly, cells expressing the CD71tm form of Tim-3 displayed significantly smaller immune synapses, as revealed by CD3 localization, suggesting a dominant inhibitory effect of this construct.

**Figure 5.**
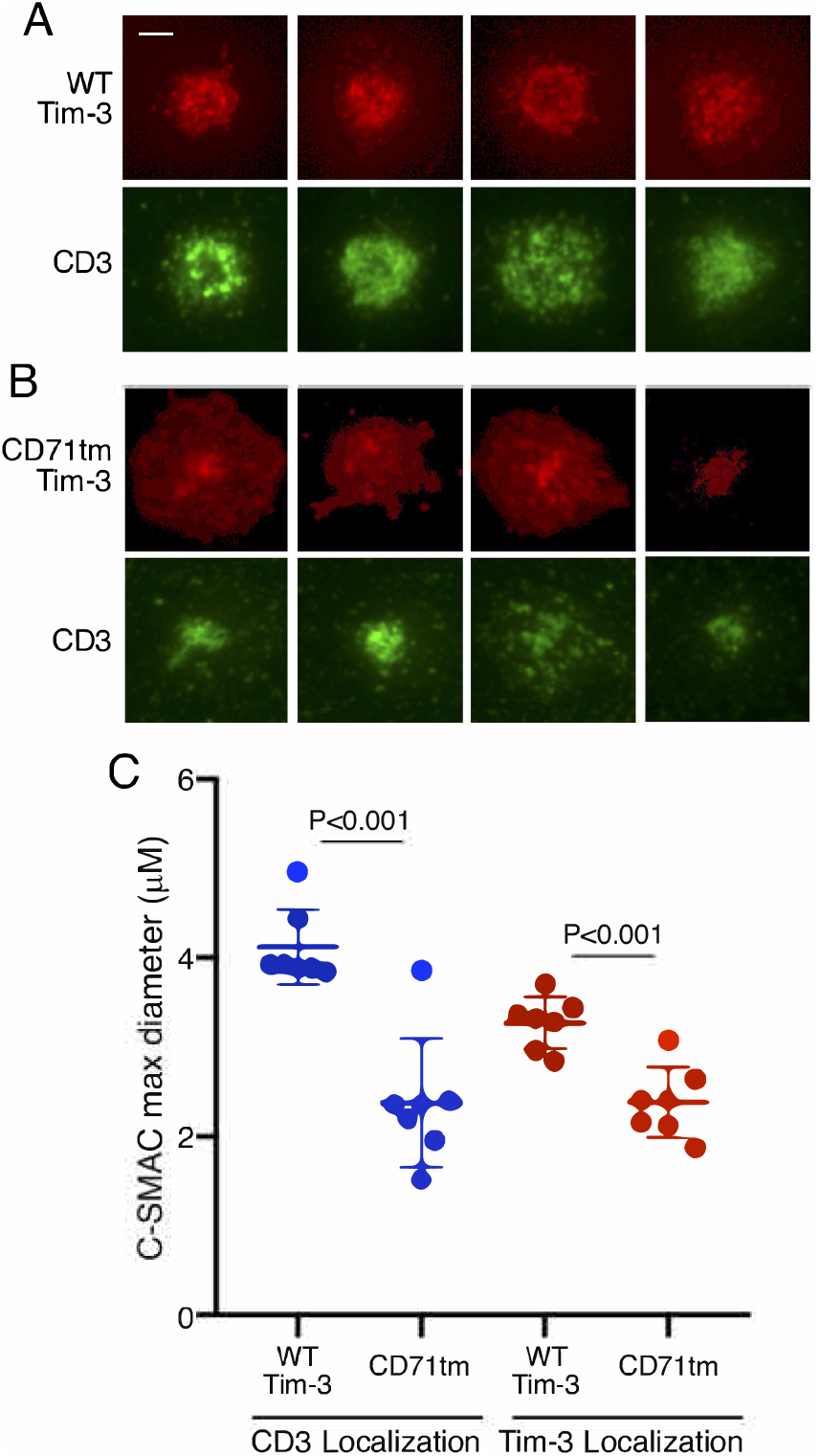
Lipid bilayer imaging of immune synapses with WT vs. CD71tm Tim-3. **(A)**Representative images of murine CD8^+^ T cells expressing FAP-tagged WT Tim-3 (top rows) or CD71tm Tim-3 (bottom rows). Tim-3 localization was determined based on FAP staining after addition of dye. TCR/CD3 localization is tracked by a labeled anti-TCR (H57) Fab fragment (green). Images were acquired at 15 mins. after addition of T cells to the bilayer. The scale bar is 2μM. (**B**) Quantitation of average Tim-3 and CD3 synapse diameter from 50-60 cells (Tim-3) or 80-90 cells (CD3). Statistical significance was derived from student’s t tests of WT vs. CD71tm.

### Enforced IS localization of the Tim-3 cytoplasmic tail is still permissive for T cell activation by a chimeric antigen receptor

As discussed above, multiple ligands have been reported to bind to the IgV domain of Tim-3, although none of these is exclusive to Tim-3. The preceding experiments here were conducted in the absence of added ligands to the ecto domain of Tim-3, although we cannot rule out the presence of such ligands in our cultures. We therefore considered the possibility that ligand engagement of Tim-3 could affect its ability to localize to the IS and/or provide a co-stimulatory signal. While several ligands for Tim-3 have been described (*8, 49*), including galectin-9, HMGB1, PS and CEACAM1, none of these is exclusive to Tim-3. We therefore tested the effects of enforced localization of Tim-3 to the IS, using a chimeric antigen receptor (CAR) approach. To this end, we compared the activity of an anti-CD20 CAR containing the cytoplasmic domains of TCR ζ and CD28 (*50, 51*) with one containing the cytoplasmic domains of ζ Tim-3. Initially, we expressed these receptors in Jurkat T cells, which were cultured with CD20-expressing Raji B cells to generate conjugates, as in the above experiments. As expected, both of these CARs formed discrete IS’s when bound to cells expressing cognate antigen (**Fig. 6A**).

**Figure 6.**
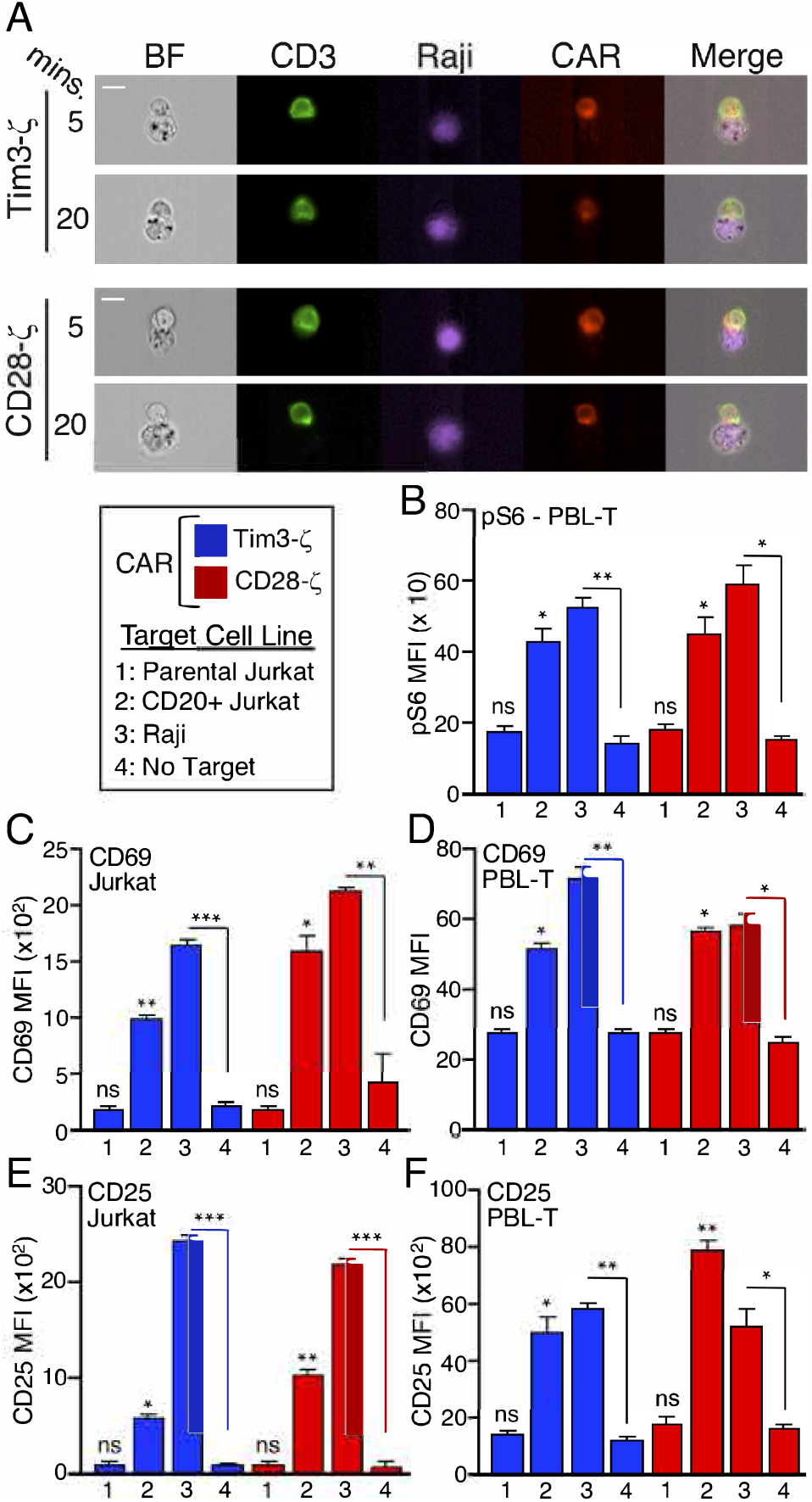
Enforced immune synapse localization of the human Tim-3 cytoplasmic tail supports T cell activation through a chimeric antigen receptor. (**A**) ImageStream analysis of immune synapse formation by the CD28-ζ or Tim3-ζ CAR, expressed in Jurkat T cells and mixed with Raji B cells. Each bar represents an average of 450 conjugates. (**B**) Peripheral blood T cells expressing either Tim3-ζ or CD28-ζ CAR were assayed by intracellular flow cytometry for phosphorylation of ribosomal protein S6 after stimulation for 30mins with the indicated target cells. (**C-F**) Jurkat T cells (**C,E**) or human peripheral blood T cells (**D,F**) expressing either the CD28 or Tim-3 CAR were stimulated with the indicated target cells (see key for panel B) and analyzed 18 hrs. later by flow cytometry for cell-surface expression of CD69 (**C-D**) or CD25 (**E-F**). Data shown represent the mean ± SEM. \**P* < 0.05, \*\**P* < 0.01, \*\*\**P* < 0.001, Tukey’s multiple comparisons test, based on three independent experiments. All statistics indicate comparison with no-target conditions. Data are representative of two experiments. The scale bars in all panels indicate 10μM.

We next assessed the functional consequences of this enforced localization of the Tim-3 cytoplasmic tail to the IS. We mixed Jurkat T cells expressing either the CD28- or Tim3-containing CAR with Raji B cells (CD20+) as targets. Given our previous findings (above and (*22, 24*)) that Tim-3 is capable of co-stimulating TCR-mediated phosphorylation of the ribosomal S6 protein, we determined whether this activity is retained in the Tim-3 CAR. Indeed, consistent with our functional data, the Tim-3 CAR mediated robust pS6 upregulation, similar to the function of the CD28 CAR (**Fig. 6B**). This was observed with two different target cell lines expressing CD20 – either Raji cells (which express endogenous CD20) or Jurkat T cells transfected with CD20. Using this approach, we also found that the Tim3-containing CAR was roughly as efficient as a CD28-containing CAR at mediating upregulation of the early activation marker CD69 when the CARs were expressed in either Jurkat or primary human T cells (**Fig. 6C**). This activity was dependent on the presence of the CD20 antigen on target cells, since it was not observed in the absence of target cells or in the presence of parental Jurkat T cells, which do not express CD20. We also assessed the effects of the two CARs on induced expression of CD25, expression of which is increased with somewhat delayed kinetics, relative to CD69. Consistent with the effects shown above, the CD28 and Tim-3 CARs mediated antigen-specific upregulation of CD25 to a similar extent, in both Jurkat and primary human T cells (**Fig. 6D**). Thus, in the context of a synthetic antigen receptor, which eliminates potential confounding effects of Tim-3 ligands, the cytoplasmic tail of Tim-3 is still permissive for robust T cell activation, similar to a standard CD28-containing CAR.

## Discussion

Despite studies linking expression of Tim-3 to T cell dysfunction, molecular mechanisms underlying the biological effects of Tim-3 remain unresolved. In particular, mechanisms by which Tim-3 might directly inhibit TCR or co-stimulatory signaling are not well-understood. Work presented here is consistent with previous reports from our lab and others that Tim-3 can paradoxically mediate positive, co-stimulatory effects on T cell activation, at least under some circumstances. These effects are tightly correlated with enhanced phosphorylation of the ribosomal S6 protein. Since S6 phosphorylation can be increased by either MEK/ERK or Akt/mTOR signaling, we used specific inhibitors to probe the roles of these specific pathways in the co-stimulatory effect of Tim-3 on S6. Our data suggest that both of these pathways are involve in Tim3-mediated co-stimulation. This is consistent with previous findings from our group showing that Tim-3 can also enhance TCR-proximal signaling at the level of PLC-γ1 phosphorylation (*24*). Interestingly, we also found that expression of Tim-3 itself appears to be sufficient for at least partial upregulation of ERK/AP-1 activation, something that we have not observed with other downstream pathways like pAkt, pS6, NFAT, which all require additional stimulation through TCR/CD28.

Proper localization of numerous proteins during T cell recognition of antigen/APC is important for early T cell activation. For example, a PKC θ mutant that fails to accumulate at the IS impairs T cell activation, even upon stimulation with α-CD3 and α-CD28 mAbs (*52*). In agreement with a previous report from Ostrowski and colleagues with human PBL T cells (*34*), we observed stimulation-dependent recruitment of Tim-3 to the IS on both Jurkat and primary murine T cells engaged by antigen/APC. Here we extended these findings by examining the role of specific domains within Tim-3 for this recruitment, in parallel with experiments quantifying the effects on downstream signaling. Surprisingly, recruitment of Tim-3 to the immune synapse was not dependent on the ecto or intracellular domains of Tim-3, but rather appeared to be primarily determined by the transmembrane (TM) domain. Thus, previous data demonstrated TM domain-dependent recruitment of CD148 to lipid rafts, which could be disrupted by replacing the CD148 TM domain with that of CD71 (*53*). Since Tim-3 IS recruitment was previously found to be associated with lipid raft recruitment (*34*), we replaced the TM domain of Tim-3 with that of CD71. While this did not affect cell-surface expression of Tim-3, it did significantly impair both the recruitment of Tim-3 to the synapse and the ability of Tim-3 to co-stimulate TCR-mediated increases in pS6.

One of the unresolved aspects of Tim-3 biology concerns the roles of the multiple ligands that have been reported to interact with the ecto domain of the protein. Thus, galectin-9, HMGB1, phosphatidylserine (PS) and CEACAM-1 have all been reported as Tim-3 ligands (*8, 9, 49*). However, none of these is exclusive to Tim-3, as all of these putative ligands can also interact with other cell-surface receptors (*9, 17*). We cannot assume that none of the Tim-3 ligands is present in our experiments, since the APC’s used for synapse formation may express one or more of them. Relevant for this point, a Tim-3 construct lacking the IgV domain was still found in the synapse at the same level as WT Tim-3. Thus, our results suggest that ligand binding is not required for localization of Tim-3 in the synapse, since all known ligands bind to Tim-3 through its IgV domain.

Lipid raft-associated proteins are known to be recruited into the IS (*54*). Of note, several transmembrane proteins with immune function have been shown to partition into lipid rafts in a manner dependent on their specific transmembrane domains (*55–57*). Thus, when we replaced the TM domain of Tim-3 with one previously shown to be excluded from lipid rafts (CD71; (*46*)), we observed significantly decreased synapse recruitment of Tim-3. It is still not clear how Tim-3 would partition into lipid rafts, as it is not known to be acylated. However, the TM domain does contain a GxxxG motif, which was previously shown to form a helical structure that can mediate not only helix-helix association but also interaction with membrane cholesterol in the case of amyloid precursor protein (APP) (*58*). Curiously, a similar domain can be found in the TM domain of CD71. However, the context (i.e. surrounding sequence) is distinct between the TM domains of Tim-3 and CD71 and, based on previous work, this could influence the differential effects of these domains on synapse recruitment (*59*). An unexpected aspect of the CD71 TM Tim-3 construct was that it was associated with less robust synapse recruitment of CD3. We do not yet know whether this was a direct effect or rather just a reflection of relatively lower synapse recruitment in comparison to cells ectopically expressing Tim-3. Indeed, we previously reported that Tim-3 could recruit a significant fraction of Fyn to its cytoplasmic tail (*24*) and another study suggested a role for Fyn recruitment in stabilizing TCR synapses (*60*). Thus, it is possible that non-synapse localized Tim-3 with a CD71 TM domain is sequestering Fyn away from active synapses. Nonetheless, this is not sufficient to completely shut down CD3 signaling, since anti-CD3 induced activation of pS6 was not affected by the CD71 TM mutant.

Returning to the question of stimulatory vs. inhibitory signaling by Tim-3, results presented here are consistent with previous studies from our lab (*22, 24*) and others (*23, 61, 62*) suggesting a co-stimulatory role for Tim-3 in T cells. As suggested by others, interactions of Tim-3 *in trans* with other transmembrane proteins like CEACAM1 (*42*) or phosphatases (*34*) could modulate the signaling function of Tim-3 toward an inhibitory pathway. Nonetheless, some of our cytoplasmic tail deletion mutants (T2 and Δcyto constructs) lose their co-stimulatory activity, while still localizing to the synapse. Perhaps more strikingly, even enforced localization of Tim-3 to the synapse through the use of a CAR still allowed for efficient T cell activation upon stimulation with cells expressing the ligand for the CAR. This provides strong support for our conclusion that the cytoplasmic tail of Tim-3 contains intrinsic co-stimulatory activity.

Most literature on Tim-3 in T cells has focused on exhausted T cells and endogenous Tim-3 is not expressed to any detectable degree by naïve T cells in either mice or humans. Thus, how do our findings relate to the *in vivo* function of Tim-3? We previously speculated that enhanced TCR signaling by Tim-3 may be important for the acquisition of T cell exhaustion, a process known to be tightly linked to chronic stimulation (*27*). Based on our recent work (*22*) and that of Colgan and colleagues (*62*), Tim-3 is also transiently upregulated under conditions of acute T cell activation, so the protein may be playing a role during the effector phase. Similarly, we also found that Tim-3 is maintained, albeit at a low level, on memory T cells, where it may play a role in re-activation of these cells (*22*). Tim-3 is constitutively expressed by mast cells and some cells of the monocytic lineage (e.g. CD103+ DCs). Thus, the signaling pathways downstream of Tim-3 that we have described here and elsewhere are potentially also of relevance for those cells, including under steady-state conditions.

Although much of the evidence is indirect, many studies in the literature have concluded that Tim-3 is an inhibitory molecule. Going forward, it will be important to reconcile findings of positive effects of Tim-3 signaling with previous reports of Tim-3 inhibitory function within the T cell compartment. This is an important issue, since antibody therapies directed at Tim-3 are being developed, with some already in clinical trials, for immune checkpoint blockade. In addition, our finding that the cytoplasmic tail of Tim-3 still allows for positive function of a CAR opens the way for more detailed study of the possible therapeutic benefits of CAR-T cells expressing such constructs.

## Materials and Methods

### Cell Lines

Jurkat and D10 T cell and Raji B cell lines were maintained in RPMI media supplemented with 10% bovine growth serum (BGS; Hyclone), penicillin, streptomycin, L-glutamine, non-essential amino acid, sodium pyruvate, HEPES and 2-mercaptoethanol. Recombinant human IL-2 (25 U/ml) was also supplemented for D10 cells. Human embryonic kidney (HEK) 293 cells were maintained in Dulbecco’s MEM supplemented with 10% bovine growth serum (BGS), penicillin, streptomycin and L-glutamine.

### Mice

FSF-Tim3 knock-in mice were generated as previously described (*22*). P14 TCR transgenic mice were obtained from Jackson Laboratories. E8i-Cre mice were obtained from Dario Vignali, and were originally generated by Taniuchi and colleagues (*63*). WT C57BL/6 mice were either obtained from Jackson Laboratories or bred in-house. Mice were age-matched within experiments. Animals were maintained in facilities of the University of Pittsburgh Division of Laboratory Animal Resources. All animal studies were performed in accordance with University of Pittsburgh Institutional Animal Care and Use Committee guidelines.

### Transfections and activation

Jurkat and D10 T cells were resuspended in RPMI media without supplements and electroporated with control plasmid, Flag-tagged WT Tim-3 or Flag-tagged mutant Tim-3 constructs, along with pMaxGFP plasmid in a Bio-Rad GenePulser at 260V and 960μF for Jurkat T cells and 250V and 950μF for D10 cells.

Transfected cells were starved of serum in phosphate-buffered saline (PBS) with 0.1% bovine serum albumin (BSA) for 1hr at 37°C. Cells were also pretreated with U0126 (Millipore Sigma #662005) and/or Akti1/2 (Abcam #ab142088) before being stimulated with the anti-Jurkat TCR Vβ8 monoclonal antibody C305. After stimulation, cells were washed in staining buffer (1% BGS-supplemented PBS) and incubated with anti-Flag-PE (clone L5; BioLegend #637310) and Ghost Dye Violet 510 (Tonbo Biosciences #13-0870-T100) at 4 ̊C for extracellular staining. Cells were then fixed and permeablized with the BD Cytofix/Cytoperm kit (BD Biosciences #554714). Fixed and permeablized cells were then incubated at 4 ̊C with anti-phospho-S6 (Ser^235/236^) APC (clone D57.2.2E; Cell Signaling Technology #14733). Samples were acquired on a BD LSRII flow cytometer, and data were analyzed with FlowJo software.

### Luciferase assays

Jurkat T cells were transfected by electroporation as described immediately above, with empty vector (pCDEF3) or various Tim-3 constructs, plus luciferase reporter driven by multimerized NFAT or AP-1 elements. The next day, cells (10^5^/well) were cultured in 96-well U-bottom plates for six hours with the indicated stimuli, after which plates were then placed at −80°. Luciferase analysis was performed as previously described (*64*). Briefly, plates were thawed at room temperature, followed by addition of lysis buffer containing NP-40 and assay buffer containing Mg/ATP. Luciferase activity was determined in a 96-well luminometer (Promega) with injection of luficerin.

### Western blotting

Cells were lysed in ice-cold NP-40 lysis buffer (1% NP-40, 1 mM EDTA, 20 mM tris-HCL pH 7.4, 150 mM NaCl) with protease inhibitors. Proteins were separated by 10% SDS-polyacrylamide gel electrophoresis and were transferred onto polyvinylidene difluoride membranes which were then blocked in 4% BSA. The membranes were then incubated with the primary antibodies overnight. This was followed by incubating the membrane with HRP-conjugated secondary antibodies for 2hrs before detection with the SuperSignal West Pico ECL substrate (Thermo Fisher Scientific #34577) and imaging on a Protein Simple FluorChem M.

The following antibodies were used for western blotting: Direct-Blot HRP anti-phospho-ERK (Thr^202^/Tyr^204^) (clone 4B11B69) - BioLegend #675505; anti-Erk1/2 - Cell Signaling Technology #9102; anti-phospho-Akt (Thr^308^; clone 244F9) - Cell Signaling Technology #4056; anti-Akt (clone 2, PKBα/Akt) -BD Biosciences #610877.

### IP’s and glycosidase treatment

HEK 293T cells (10^6^) were transfected in 6-well plates with corresponding plasmids with Transit LT1 transfection reagent (Mirus) following the manufacturer’s protocol. Transfection efficiency was determined by flow cytometry. Transfected cells (5×10^5^ per well) were lysed using for one hour with ice-cold 1% NP-40 lysis buffer (20 mM Tris-HCl, pH 7.5, 150 mM NaCl, 1 mM EDTA) supplemented with complete mini (Roche) and HALT (Invitrogen) protease inhibitors.

Protein-G agarose beads were washed and blocked with 1% BSA for 30 min. Whole cell lysate and beads were incubated with 1 μg anti-Flag antibody M2, rotating at 4° C overnight. Beads were pelletted by centrifugation and washed 3x with lysis buffer for or with deglycosylation buffer. Deglycosylation was performed on beads using Protein Deglycosylation Mix II (NEB) in denaturing conditions according to manufacturer’s protocol. Beads were boiled with 2x Sample buffer (Bio-Rad) with β-ME for 5 minutes at 95 degrees. Samples were spun down and protein supernatant was removed. Supernatant was run on 10% Nupage gels and transferred to PVDF membrane. Membranes were blocked with 5% BSA in TBST buffer and probed with anti-mouse Tim-3 goat polyclonal antibody (R&D) and detected with anti-goat-HRP antibody and SuperSignal West Pico Chemiluminescent reagent (Thermo Fisher).

### Jurkat-Raji conjugate formation

Raji cells were labeled with either CellTracker Orange CMRA Dye (Thermo Fisher Scientific #C34551) or Cell Proliferation Dye eFluor450 (Thermo Fisher Scientific #65-0842-85) and then pulsed with 2 μg/ml staphylococcal enterotoxin E (SEE) (Toxin Technology #ET404) for 1hr at 37°C. Transfected Jurkat T cells were mixed with Raji cells, either unpulsed or pulsed with SEE, at a ratio of 1:1 and incubated for indicated times at 37°C.

### Primary T-B cell conjugate formation

B cells from spleens and lymph node of naïve mice were purified by magnetic separation using a pan-B cell isolation kit (Miltenyi Biotec #130-095-813), then incubated overnight with 30 μg/ml LPS (Sigma #L2630) and 10 μM LCMV gp33-41 peptide (AnaSpec #AS-61296). CD8 T cells from spleens and lymph nodes of either P14 or FSF-Tim-3/E8i-Cre/P14 mice were purified by magnetic separation using a naïve CD8^+^ T cell isolation kit (Miltenyi Biotec #130-104-075) and then mixed with gp33-pulsed B cells at a ratio of 1:1 and incubated for indicated times at 37°C.

### Imaging flow cytometry

Cell conjugates were fixed with 2% paraformaldehyde and then incubated with antibodies at 4 ̊C for extracellular staining. For intracellular staining, cells were permeabilized with 0.2% saponin and then incubated with anti-phospho-S6 (Ser^235/236^) antibody at 4 ̊C.

Imaging cytometry data on fixed and stained cells were acquired on an Amnis ImageStreamX Mark II running INSPIRE^®^ software (Millipore Sigma). Initially, 10^4^ events were acquired, before downstream gating and analysis, which was performed using IDEAS^®^ software (Millipore Sigma). First, focused cells, based on the gradient root mean square feature, were identified. Doublets were identified by the aspect ratio and area of bright field and then gated by the aspect ratio and staining intensity of B cells. Gated cells were further refined by the aspect ratio and staining intensity of T cells to identify T-B cell doublets. For Jurkat T-Raji conjugates, an interface mask covering the immunological synapse was made from brightfield and Raji staining. The percentages of CD3 and Tim-3 recruitment were quantified from the ratio of the intensity of either CD3 or Tim-3 in the interface mask to that of the whole cell mask. The intensity of pS6 in Jurkat T cells was quantified in the mask made by subtracting the B cell mask from the brightfield mask. For primary T-B conjugates, an interface mask was made from brightfield and CD8 staining. The intensity of pS6 in CD8+ T cells was quantified in the mask made from CD3 staining. Approximately 100–300 cells per point are represented in the Jurkat T-Raji conjugate data, and 100–2,000 cells are represented in the primary T-B conjugate data.

The following antibodies were used for conjugation assay: anti-human CD3-FITC (clone OKT3; Tonbo Biosciences# 35-0037-T100), anti-Flag-APC (clone L5; BioLegend #637308), anti-mouse CD3-FITC (clone 145-2C11; Tonbo Biosciences #35-0031-U500), anti-mouse CD19-violetFluor450 (clone 1D3; Tonbo Biosciences #75-0193-U100), anti-pS6 (Ser^235/236^)-PE (clone D57.2.2E; Cell Signaling Technology #5316S).

### TIRF imaging of T cell synapses on lipid bilayers

WT and CD71tm mutant Tim-3 constructs were cloned into MSCO-based retroviral vectors containing the fluorogen activating protein (FAP) sequence in-frame, followed by an IRES and coding sequence of mouse Thy1.1. These constructs were then transfected into 293T cells, along with the pCL-Eco packaging plasmid, using TransIt transfection reagent (Mirus).

CD8^+^ T cells were purified from C57BL/6 mice using a mouse CD8^+^ T cell isolation kit from Miltenyi. Cells were stimulated for 27 hrs on plates coated with anti-CD3/CD28 mAbs, plus 100 U/ml recombinant human IL-2. Activated T cells were harvested and transduced with FAP-encoding virus from the 293T cell transfectants. Transduced Thy1.1^+^ cells were purified by sorting and kept on ice until imaging.

Lipid bilayers were constructed as described (*65*), with some modifications (*66*). Briefly, liposomes comprised of 90% dioleoylphosphocholine, 10% DOGS (1,2-dioleoyl-SN-Glycero-3-{[N(5-amino-1carboxypentyl) iminodiacetic acid]succinyl}), and 0.2% biotin-CAP-PE (Avanti Polar Lipids) were deposited on glass coverslips cleaned with piranha solution (50:50 mixture of 30% H2O2 and 96% H2SO4). Alexa-488 streptavidin, Alexa-647 streptavidin or unconjugated streptavidin and biotinylated TCRβ mAb were sequentially loaded onto the bilayer, while unlabeled poly-his-tagged ICAM1 produced in a baculovirus system was loaded to facilitate T cell adhesion.

Transduced T cells were sorted, rested 24-48 hrs, incubated with MGnBu (200nM) in RPMI without serum or phenyl red for 15 mins. at 37°C, washed, resuspended in RPMI without serum or phenyl red and stimulated on a planar lipid bilayer. Imaging was performed using a Nikon Ti inverted microscope equipped with motorized TIRF arm and 100x 1.40 N.A. objective. Images were acquired at 100ms intervals continuously for up to 15-20 mins with a Zyla sCMOS camera (Andor) equipped with bandpass emission filters for DAPI, FITC, TRITC, and Cy5 spectral profiles.

Images were analyzed using the NIS elements software version 5.21.00. Images were separated by layer and threshold for each layer was set. AF488 threshold was set between 200-1000, AF647 threshold was set between 150-40,000. Areas for individual channels and intersecting areas were measured using automated measurements.

### Construction and characterization of chimeric antigen receptors

To generate the TIM-3 CAR constructs (*pHR-EF1-aCD19-TIM3ζ-T2A-TagBFP & pHR-EF1-aCD20-TIM3ζ-T2A-TagBFP*), DNA encoding the human TIM-3 cytoplasmic domain was codon optimized, synthesized (Integrated DNA Technologies), and cloned into the pHR-EF1-aCD19-28ζ or pHR-EF1-αCD20-28ζ CAR lentiviral expression vectors, respectively, to replace the CD28 co-signaling domain using isothermal assembly.

To generate virus, HEK293T cells (ATCC) were transfected with pVSV-G (VSV glycoprotein expression plasmid), pMD2.G and the CAR-expression plasmid by calcium phosphate transfection (Clontech). At 16 hrs post-transfection cells were washed with PBS and incubated in complete DMEM media containing 6 mM sodium butyrate (Sigma Aldrich). Supernatants were collected at 24 and 48 hrs and were subsequently combined and filtered through a 45 μm vacuum filter. Viral particles were concentrated by ultracentrifugation for 1.5 hrs at 24,500 rpm, and viral pellets were re-suspended in 0.05 ml of supplemented RPMI medium and frozen at −80°C.

For CAR-T pS6 analysis, 100,000 Jurkat or primary T cells expressing the TIM-3 or CD28 CAR were incubated with an equal number of the indicated antigen positive or negative target cell line in a V-bottom plate for 30mins at 37°C in 5% CO2. Cells then underwent intracellular staining and flow cytometry as described (*22*).

CAR-T surface marker analysis: 100,000 Jurkat or Primary T cells expressing the TIM-3 or CD28 CAR were incubated with an equal number of the indicated antigen positive or negative target cell line in a V-bottom plate for 18 hrs at 37°C in 5% CO2. Cells then underwent cell-surface antibody staining and flow cytometry (*22*).

## Supplementary Materials

Fig. S1. Effects of MEK and Akt inhibitors on induction of pS6 by Tim-3 and effects of transgenic Tim-3 on T cell proliferation.

Fig. S2. Expression of Tim-3 cytoplasmic truncations and their effects on pS6 induction.

Fig. S3. Retroviral system used for TIRF imaging of immune synapses.

## Acknowledgements

We thank the University of Pittsburgh Department of Immunology flow cytometry core and Center for Biologic Imaging for valuable technical assistance. JL also acknowledges the support of Dr. Olivera J. Finn.

## Author contributions

SK performed and analyzed ImageStream experiments, signaling experiments and wrote the initial draft of the manuscript. PM performed retroviral infections followed by TIRF imaging and analysis, as well as flow cytometry quantitation of Tim-3 construct expression, western blotting +/− glycosylation and Tim-3 Tg T cell pERK analysis. CJW generated lipid bilayers and performed TIRF imaging. HB and ASW performed experiments. JL provided conceptual input for the CAR-T approach, designed and generated the lentiviral constructs and performed CAR-T functional experiments. MK performed CAR-T experiments to assess pS6. LPK conceived of the project, obtained funding, assisted with experimental analysis and interpretation and wrote the final manuscript, with input from multiple authors.

## Data and materials availability

Full-length, truncated and CD71tm Tim-3 constructs have been deposited with Addgene and are available through them.

## Supplementary Materials

**Figure S1.**
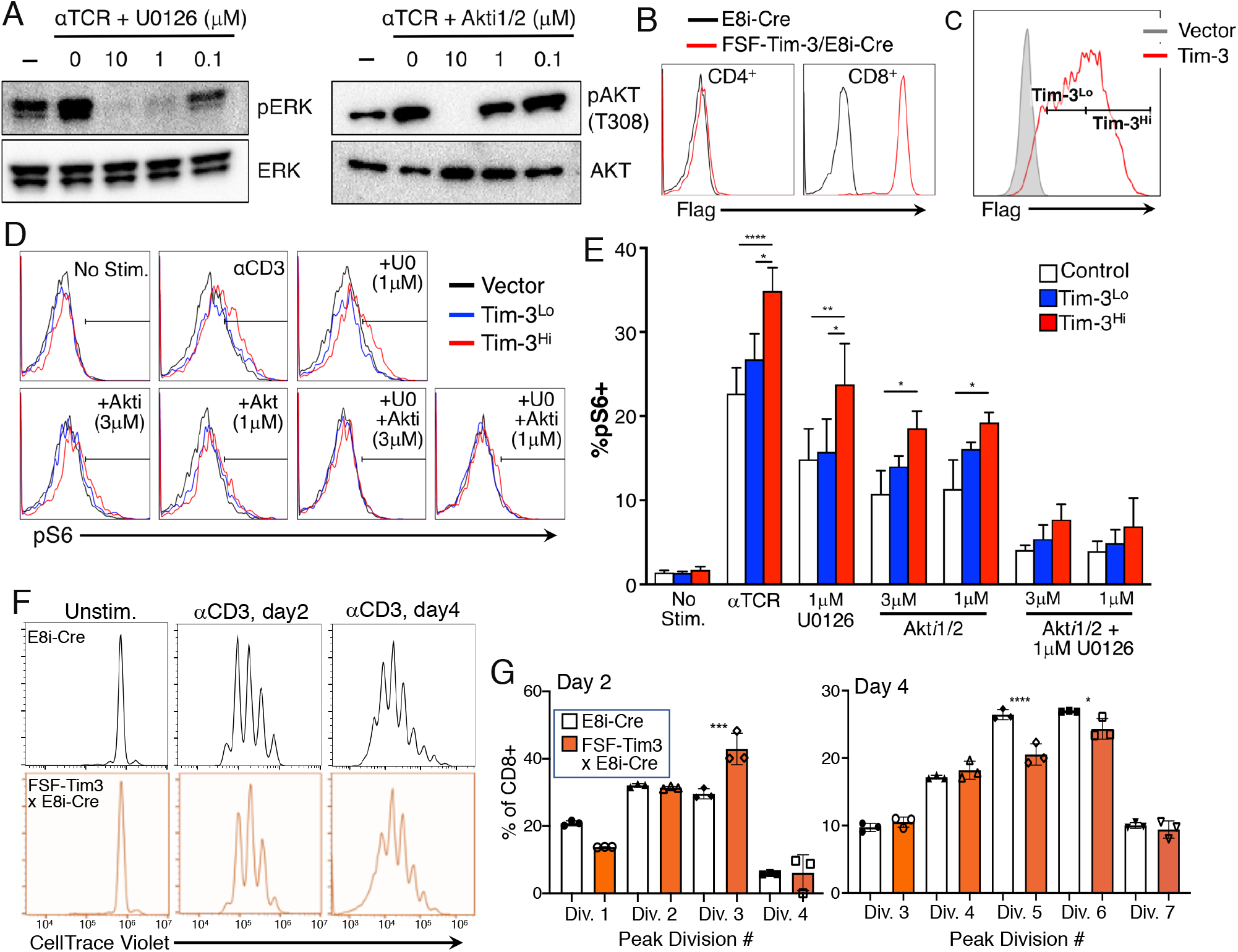
Effects of MEK and Akt inhibitors on induction of pS6. (**A**) Control experiments demonstrating the efficacy of the inhibitors on Jurkat T cells stimulated with anti-TCR mAb and the indicated concentrations of inhibitor. (**B**) Expression of Flag-Tim3 on CD4^+^ or CD8^+^ T cells from Tg mice expressing only E8i-Cre (black lines) or both E8i-Cre and the flox-stop-flox knock-in of Flag-Tim3 (red lines). (**C**) Gating used to segregate D10 T cells expressing low or high levels of Tim-3 after transfection. (**D**) Representative flow cytometry of pS6 staining of the D10 T cells analyzed in panel C, after stimulating with anti-CD3 mAb and the indicated inhibitors. (**E**) Quantitation of the fraction of pS6^+^ cells based on the flow cytometry in panel D. (**G**) Representative *in vitro* proliferation of CD8+ T cells from the indicated mice. (**H**) Quantitation of divided cells in each peak from triplicate samples. Each panel is representative of three independent experiments. Statistical significance in panels E and G was determined by one-way ANOVA.

**Figure S2.**
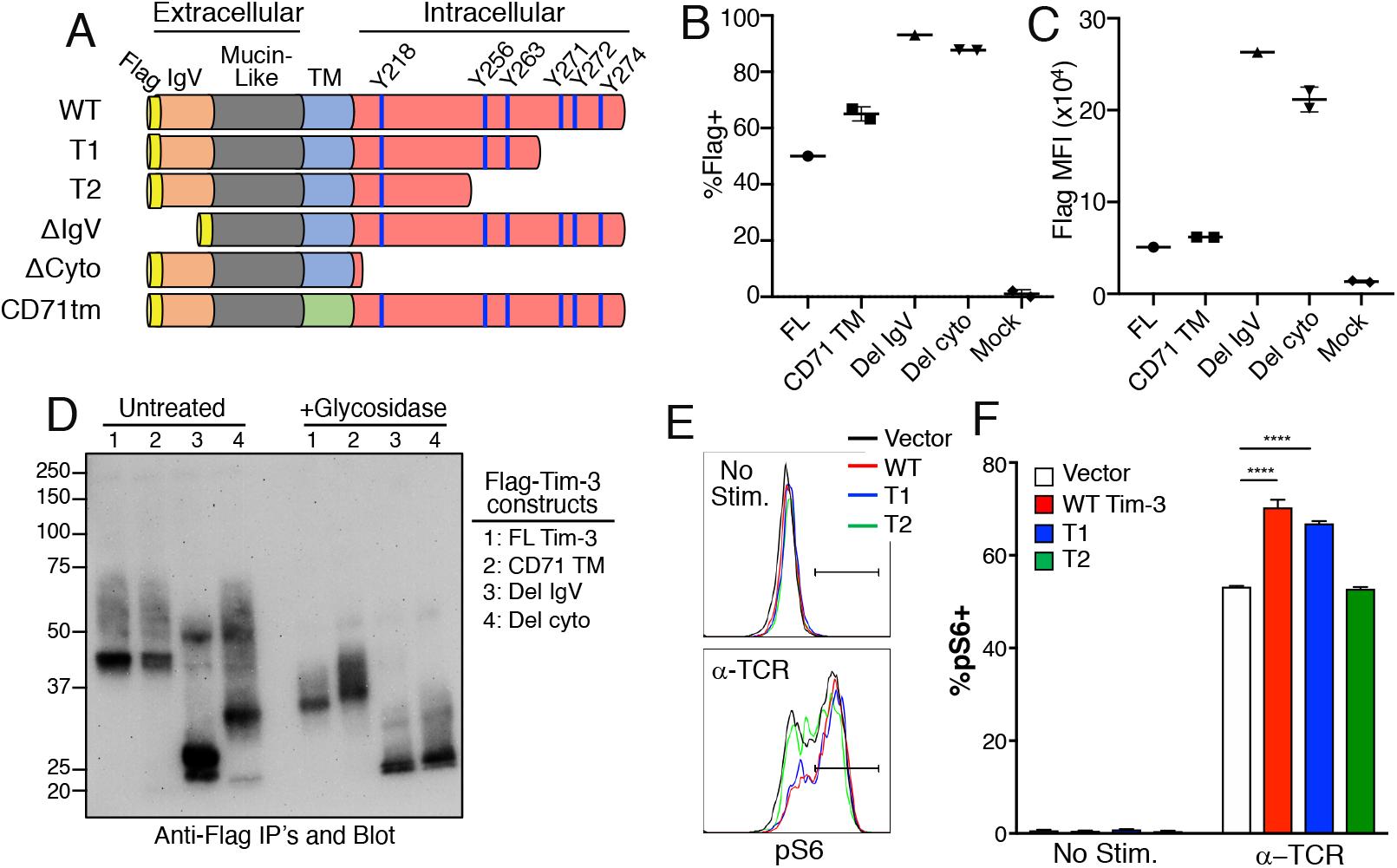
Effects of Tim-3 cytoplasmic truncations on Tim-3 expression and pS6 induction. (**A**) Schematic of the Tim-3 constructs analyzed in Figs. 3–4. (**B-C**) Cell-surface expression of Tim-3 constructs, based on flow cytometry staining for the N-terminal Flag tag. Duplicate transfections representative of three independent experiments. (**B**) Expression based on the percentage of positive cells after transient transfection of Jurkat T cells; (**C**) mean fluorescent intensity (MFI) of Flag staining. (**D**) Western blot analysis of Tim-3 constructs +/− treatment with PNGase F. Representative of three independent experiments. (**E**) Representative pS6 flow cytometry of Jurkat T cells transfected with the indicated Tim-3 constructs and stimulated with anti-TCR mAb. (**F**) Quantitation of pS6 induction in the transfected T cells. The labels “Del” and Δ refer to deletions of the indicated domains. Statistical significance in panel F was determined by one-way ANOVA.

**Figure S3.**
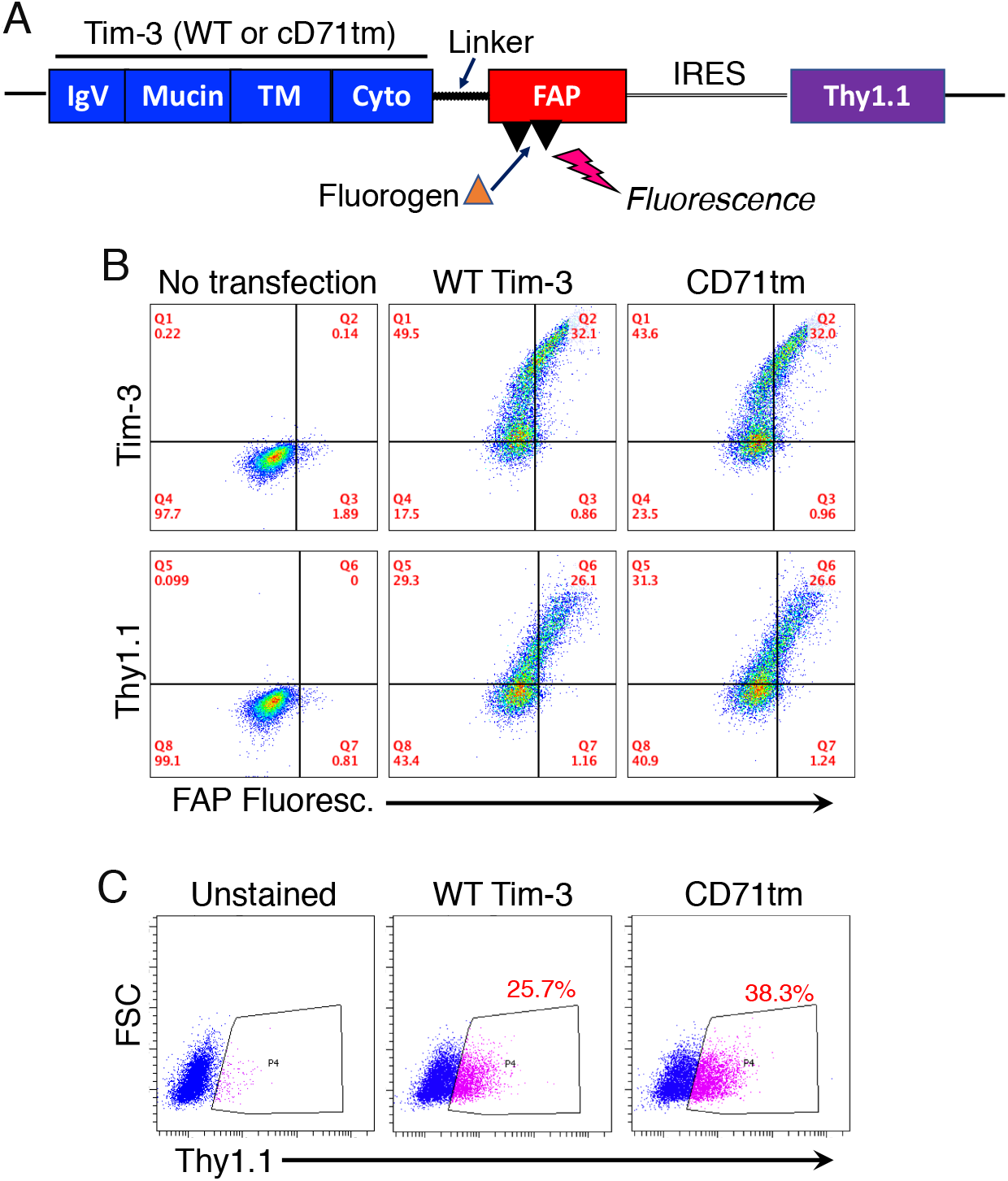
Retroviral system used for TIRF imaging of immune synapses. (**A**) Schematic of the retroviral vector used to express WT or CD71tm forms of Tim-3, along with a fluorogen-activating peptide (FAP). The vector also encodes an IRES-controlled Thy1.1 marker. (**B**) Representative flow cytometry of Tim-3 and Thy1.1 expression by the packaging (293T) cells transfected with the indicated constructs. (**C**) Representative flow cytometry of the co-encoded Thy1.1 in murine T cells transduced with the indicated retroviruses.

